# Nemacol is a Small Molecule Inhibitor of *C. elegans* Vesicular Acetylcholine Transporter with Anthelmintic Potential

**DOI:** 10.1101/2022.06.14.496146

**Authors:** Sean Harrington, Jacob Pyche, Andrew R. Burns, Tina Spalholz, Rachel J. Baker, Justin Ching, Mark Lautens, Daniel Kulke, Winnie Deuther-Conrad, Peter Brust, Peter J. Roy

**Affiliations:** Department of Pharmacology and Toxicology, University of Toronto, Toronto, ON, M5S 1A8, Canada; The Donnelly Centre for Cellular and Biomolecular Research, University of Toronto, Toronto, ON, M5S 3E1, Canada; Department of Molecular Genetics, University of Toronto, Toronto, ON, M5S 1A8, Canada; Department of Neuroradiopharmaceuticals, Institute of Radiopharmaceutical Cancer Research, Helmholtz-Zentrum Dresden-Rossendorf, 04308 Leipzig, Germany; The Department of Chemistry, University of Toronto, 80 St. George Street, M5S 3H6, Toronto, CANADA; Research Parasiticides, Bayer Animal Health GmbH, Monheim, Germany

**Keywords:** Nemacol, VAChT, nematicide, acetylcholine transporter, UNC-17, *C. elegans*, acetylcholine esterase, AChE, ivermectin, structure-activity relationship

## Abstract

Nematode parasites of humans and livestock pose a significant burden to human health, economic development, and food security. Anthelmintic drug resistance is widespread among parasites of livestock and many nematode parasites of humans lack effective treatments. Here, we present a nitrophenyl-piperazine scaffold that induces motor defects rapidly in the model nematode *Caenorhabditis elegans*. We call this scaffold Nemacol and show that it inhibits the vesicular acetylcholine transporter (VAChT), a target recognized by commercial animal and crop health groups as a viable anthelmintic target. We demonstrate that it is possible to create Nemacol analogs that maintain potent *in vivo* activity whilst lowering their affinity to the mammalian VAChT 10-fold. We also show that Nemacol synergizes with the anthelmintic ivermectin to kill *C. elegans*. Hence, Nemacol represents a promising new anthelmintic scaffold that acts through an identified viable anthelmintic target.

**One sentence summary:** A small molecule screen identifies a vesicular acetylcholine transporter inhibitor scaffold that incapacitates parasitic nematodes

## Introduction

Nematodes that parasitize humans and non-human animals including livestock are a significant burden to human health, food security and economic development. Unfortunately, most frontline anthelmintic molecules suffer from inadequacies. For example, anthelmintics used to treat human hookworm (*Ancylostoma duodenale* and *Necator americanus*) and whipworm (*Trichuris trichiura*) are recognized to have inadequate efficacy^1^. Furthermore, the top four marketed anthelmintics used in non-human animals, which include macrocyclic lactones (e.g. ivermectin), tetrahydropyrimidines (e.g. pyrantel), imidazothiazoles (e.g. levamisole) and benzimidazoles (e.g. thiabendazole), have demonstrated pervasive resistance in cattle and small-ruminants^2^ as well as companion animals^3-7^. The dire need for new anthelmintics has been recognized by academic, industry and governmental experts for some time^1,8,9^.

Towards identifying novel candidate anthelmintic scaffolds, our group has recently carried out small molecule screens for those that affect the motor activity of the free-living nematode *Caenorhabditis elegans*^10^. One scaffold that we focus on here inhibits cholinergic signalling. The neurotransmitter acetylcholine (ACh) is the primary signalling molecule in animals that triggers muscle contraction^11^. Without careful regulation of ACh, animals lose muscular control and die^12-14^. The essentiality of cholinergic signaling is a major point of vulnerability that has been repeatedly exploited, both in nature as a target of toxins and venoms^14,15^ and as a pesticide strategy^16,17^.

Three protein groups within the cholinergic pathway are targeted by anthelmintic and nematicidal molecules. These include nicotinic ACh receptors (nAChRs), which are agonized by the imidazothiazoles and tetrahydropyrimidines^3,18-21^ and antagonized by derquantel^22-24^. Acetylcholinesterase (AChE) that breaks down ACh at the neuromuscular junction is inhibited by organophosphates & carbamates, which in turn leads to catastrophically high levels of synaptic ACh. Finally, the vesicular acetylcholine transporter (VAChT), which packages ACh into presynaptic vesicles^12,25,26^, is inhibited by the relatively novel spiroindoline scaffold^27^. Spiroindolines have been investigated as nematode and/or insect parasiticides by Zoetis (patent US20160296499A1^28^), Intervet Inc (patent US9096599B2)^29^ and Syngenta (patent US9174987B2)^30^, but none have yet brought a sprioindoline to market^27^.

Here, we present a novel nitrophenyl-piperazine scaffold that inhibits VAChT. Because of its structural similarity to the VAChT inhibitor vesamicol (which is used as a tool compound^25,27^), we call this scaffold Nemacol. Nemacol rapidly induces *C. elegans* worms to coil their bodies, which is a phenotype shared by mutants of the CHA-1 acetyltransferase enzyme, which makes ACh, and mutants of the VAChT worm ortholog (UNC-17)^12,26^. Herein, we describe the kinetics of the Nemacol-induced phenotypes and provide chemical-genetic and biochemical evidence to show that Nemacol inhibits nematode VAChT. We demonstrate that Nemacol also disrupts the motor activity of the commercially important animal parasite *Dirofilaria immitis* (dog heartworm) and can synergize with the macrocyclic lactone ivermectin to kill *C. elegans*. Finally, we show that select Nemacol analogs can maintain their low micromolar potency in nematodes but reduce their affinity for the mammalian VAChT receptor 10-fold relative to the Nemacol parent, demonstrating the potential for an expanded therapeutic window. We conclude that Nemacol is a novel small molecule scaffold with anthelmintic potential.

## Results

### Nemacol Inhibits the Vesicular Acetylcholine Transporter

From our previous screen^10^, we identified four structurally similar molecules that stimulate *C. elegans* egg-laying (Egl) and elicit similar uncoordinated (Unc) motor phenotypes (Fig. 1). All four molecules share a 1-ethyl-4-(4-nitrophenyl)piperazine substructure, which we refer to here as Nemacol. Within minutes of exposure, Nemacol-1 causes jerky-Unc movement (Fig. 1a-d). Over the course of four hours, the phenotype transits to tight coiling (Supplemental Movie 1) and flaccid paralysis, and then gradually transits back to the jerky-Unc phenotype. After 24 hours of being on Nemacol-1, animals no longer have obvious motor defects (Fig. 1a). Blocking drug-metabolizing cytochrome P450s by knocking down *C. elegans* cytochrome P450 reductase (EMB-8) antagonizes the dissipation of these phenotypes (Fig. S1).

**Figure 1.**
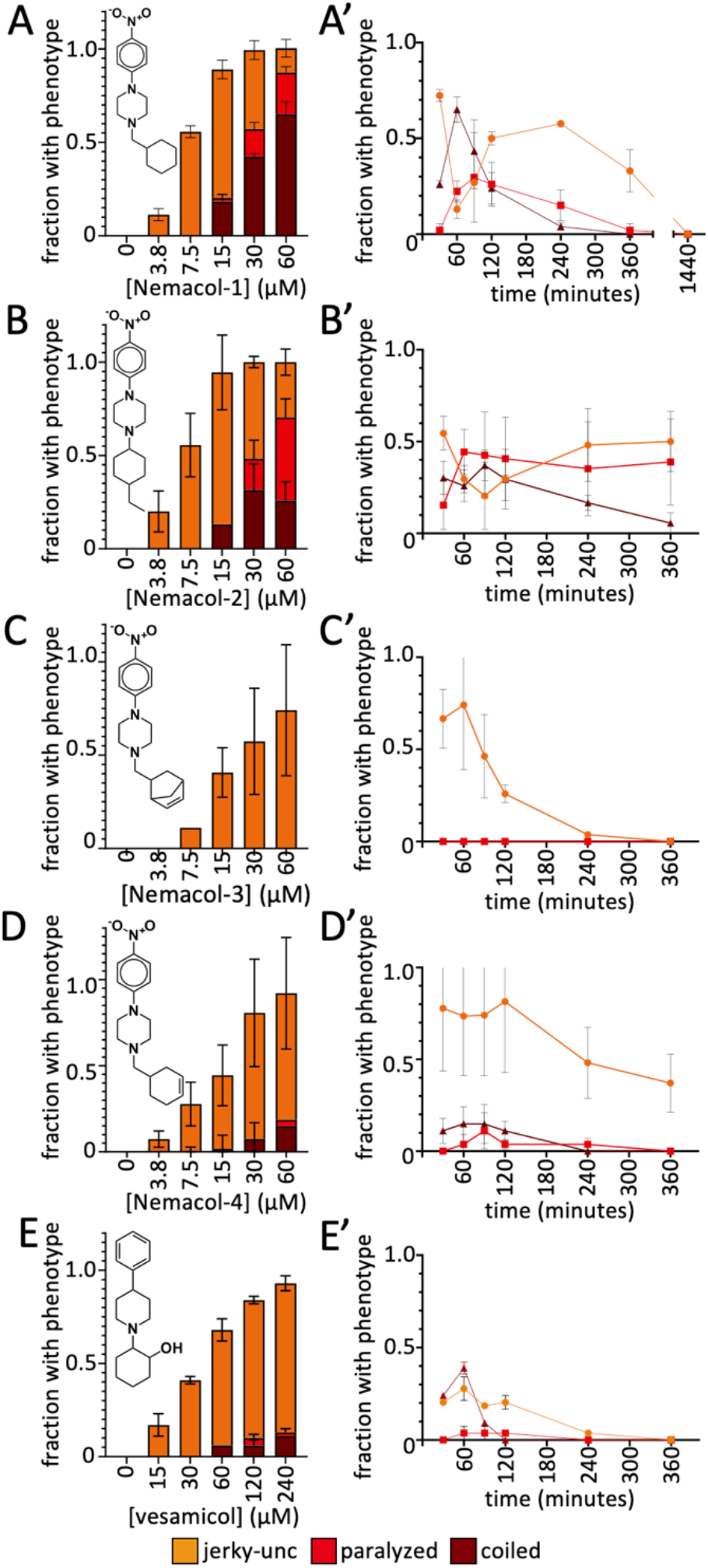
Nemacol analogs induce a shared set of phenotypes with vesamicol. (**A**) L4 wild-type worms were picked onto solid agar containing the indicated concentration at 60 minutes (left panels) and scored at the indicated time on 60 μM of indicated compound (right panels). The indicated phenotypes are scored based on subjective classification (see methods for details). Data are the mean of 3 biological replicates scoring ∼18 animals per trial showing standard deviation.

The phenotypes induced by the Nemacol scaffold are shared with *C. elegans* mutants that have defective cholinergic signaling. For example, mutations in the CHA-1 choline O-acetyltransferase that produces ACh or in the UNC-17 VAChT that packages ACh into presynaptic vesicles exhibit a characteristic coiling phenotype^12,31^. Furthermore, the Nemacol scaffold has structural similarity to the canonical VAChT inhibitor vesamicol, which induces coiling and jerky-Unc phenotypes, but with an EC50 that is 5.1-fold higher than Nemacol-1 (Fig. 1e). These similarities led to the hypothesis that Nemacol inhibits *C. elegans* UNC-17/VAChT. We tested this hypothesis in four ways.

First, we reasoned that if UNC-17 is Nemacol’s target, then weak alleles of *unc-17* (*e327* and *e795*)^26,32^ should be hypersensitive to the compound, which is what we observed (Fig. 2a).

**Figure 2.**
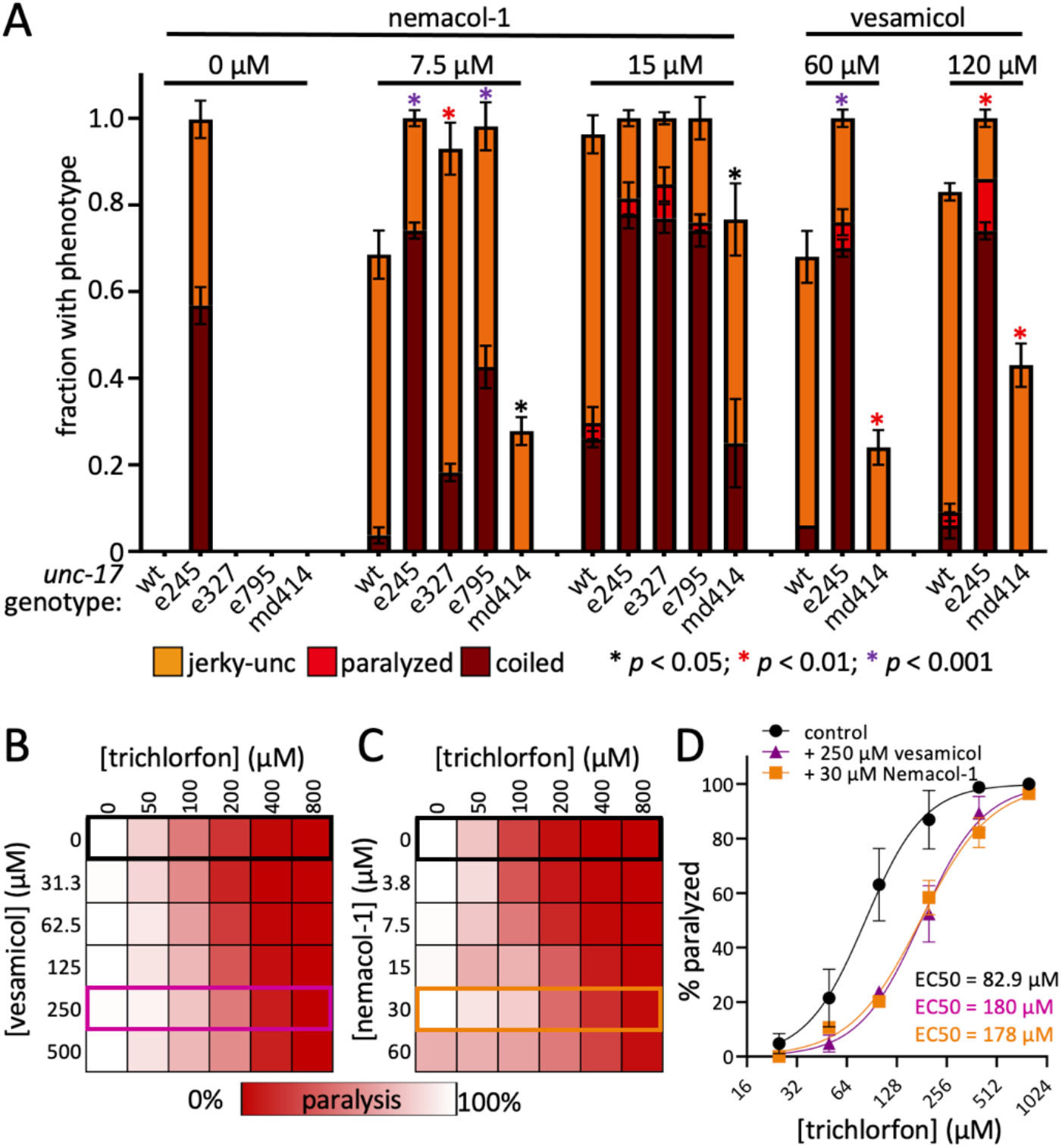
Nemacol induces phenotypes consistent with the inhibition of VAChT. (**A**) Adult wildtype worms were picked onto solid agar containing either Nemacol-1 or vesamicol at the indicated concentration at 60 minutes. The legend details are the same as that for Figure 1. Significance was calculated using Chi-square test comparing the fraction of animals exhibiting any motor phenotype between WT and the indicated test mutant. (**B-C**) Double dose-response matrices of vesamicol + trichlorfon (C) and Nemacol-1 + trichlorfon showing the fraction of animals that were scored as paralyzed after 80 minutes. The benchmark for paralysis was a lack of sinusoidal body posture and an inability to back at least half a body length upon a touch on the head with a platinum wire. Values in cells represent the % of animals scored as paralyzed. Data are the mean of 3 biological replicates scoring 28 animals per condition. (**D**) Dose-response curves from B-C highlighting the shift of trichlorfon paralysis EC50 +/- 250 μM vesamicol (purple) or 30 μM Nemacol-1. For control compared to 250 μM vesamicol *p* = 3.7 × 10^−6^; for control compared to 30 μM Nemacol-1 *p* = 5.1 × 10^−5^. *P*-value calculated using an extra sum-of-squares F test comparing the trichlorfon dose response to the indicated dose response in GraphPad Prism version 9.3.1.

Second, the *C. elegans* UNC-17 C391Y mutation called *md414* had previously been shown to disrupt vesamicol binding^32,33^. We reasoned that if Nemacol inhibits VAChT through a shared binding site with vesamicol, then *unc-17(md414)* mutant animals should suppress Nemacol phenotypes, which is what we observed (Fig. 2a).

Third, we tested whether Nemacol can suppress the paralysis induced by inhibitors of AChE. AChE inhibition results in excess ACh at neuromuscular junctions, which in turn paralyzes the worm due to excess muscle contraction^33,34^. Reduction-of-function *unc-17* mutants are known to be resistant to the effects of AChE inhibitors^26,27,33^. If Nemacol inhibits UNC-17 and results in lower levels of ACh at the synaptic cleft, then Nemacol treated worms should also resist the effects of AChE inhibitors. Indeed, Nemacol-1 can suppress the paralysis induced by two structurally distinct AChE inhibitors, namely trichlorfon (an organophosphate) and aldicarb (a carbamate) in a dose-dependent manner (Fig. 2b-c, Fig. S2). We find that 30 μM Nemacol-1 yields nearly an identical 2.2-fold shift in the EC50 of trichlorfon paralysis compared to 250 μM vesamicol representing an 8.3-fold shift in potency (Fig. 2d).

Finally, we tested whether Nemacol was able to directly interact with vertebrate VAChT. We measured the ability of Nemacol-1 to displace [^3^H]vesamicol bound to rat VAChT expressed in rat PC12^A123.7^ cells, which is an assay previously established for measuring small molecule affinity for VAChT^32,35^. We found that Nemacol-1 binds mammalian VAChT, but with 29-fold less affinity than vesamicol (Fig. 3a). Together, these data indicate that Nemacol likely elicits motor phenotypes in *C. elegans* through the inhibition of UNC-17/VAChT at the vesamicol binding site.

**Figure 3.**
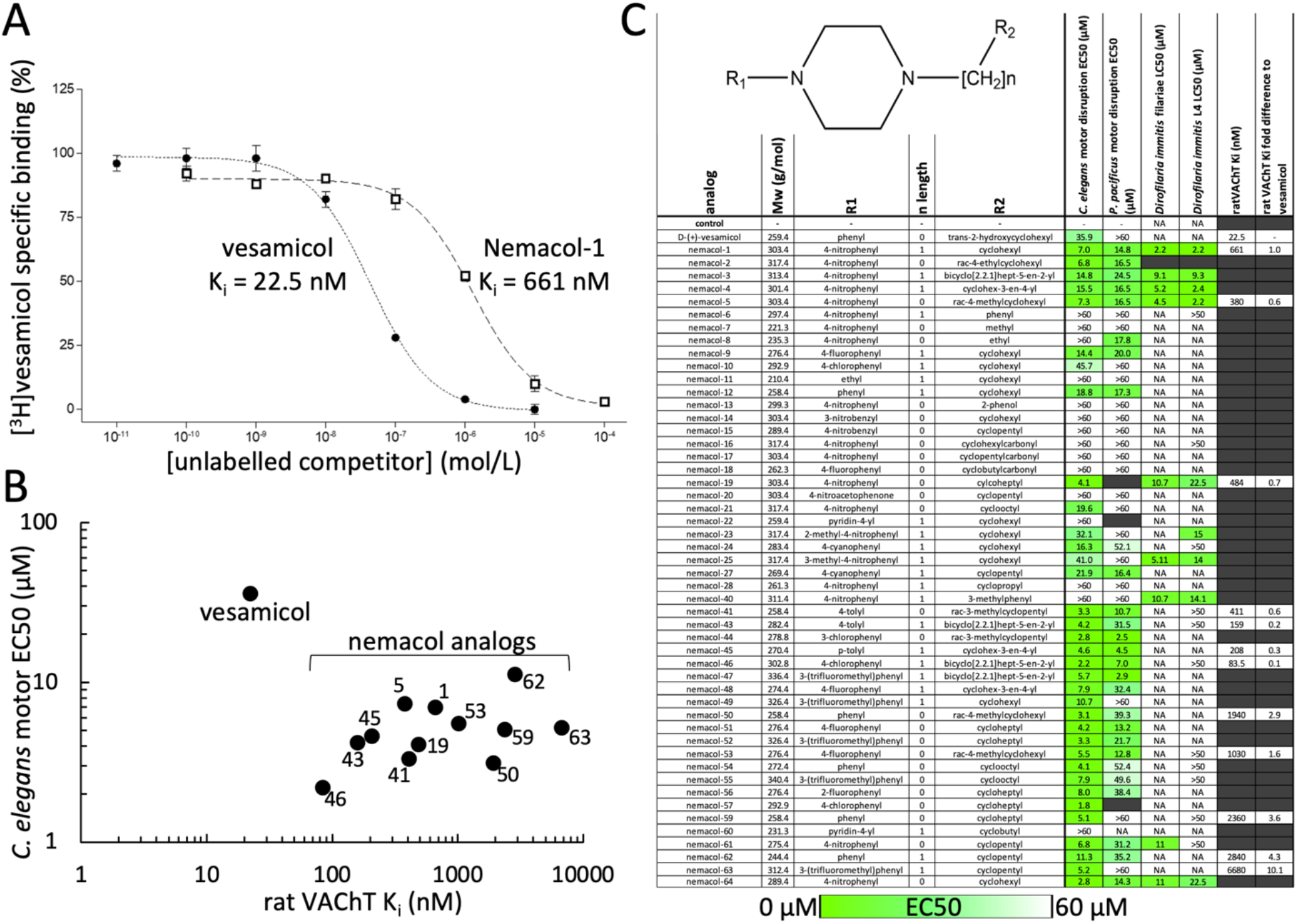
Nemacol analogs demonstrate nematode selective activity. (**A**) Nemacol-1 competitive displacement of [^3^H] vesamicol binding of rat VAChT (see methods for details). (**B**) Comparison of vesamicol and Nemacol analog *in vivo* potency and rat VAChT binding affinity. (**C**) Structure activity relationship summary of Nemacol analog activity across nematode species and rat VAChT inhibition constant (K_i_). *C. elegans* and *Pristionchus pacificus* motor assays were conducted scoring the fraction of animals demonstrating motor phenotypes on solid media containing suspended molecule as indicated. Animals were scored as ‘jerky-uncoordinated’, ‘coiled’, ‘flaccidly paralyzed’ or ‘wild-type phenotype’; data are the EC50 of animals demonstrating any motor phenotype. *Dirofilaria immitis* microfilariae EC50 data are from triplicate measurement of ∼36 animals per condition after 72h of drug exposure. *D. immitis* L3 EC50 data are from duplicate measurement of 10 freshly isolated L3s after 72 hours of incubation. The inhibition constant (K_i_) of compounds for rat VAChT, as well as the fold difference relative to vesamicol is shown (see methods for details).

### Nemacol Analogs Demonstrate Potential Selectivity Against the Nematode VAChT Target

Above, we show that Nemacol-1 is 5.1-fold more potent than vesamicol in live *C. elegans* worms, but has 29-fold less affinity than vesamicol for the mammalian VAChT target. This comparison raised the possibility that the Nemacol scaffold could be modified to create nematode-selective VAChT inhibitors. Although co-crystal structures of vesamicol with VAChT are not available, several residues known to be important for interaction are known^31,36-38^. Inspection of a VAChT multiple sequence alignment shows that residues immediately flanking vesamicol-interacting residues are divergent in nematodes (Fig. S3), raising the possibility that nematode-selectivity may be achieved.

We explored the activity of the Nemacol scaffold by testing 50 analogs in acute motor tests in culture against *C. elegans*, the free living nematode *Pristionchus pacificus*, and with select molecules, against the dog heartworm *Dirofilaria immitis*. 44 of these analogs were procured from commercial sources and 6 other analogs were synthesized by us (see methods). We found many analogs to be active against *C. elegans* and *Pristionchus*, and two analogs (Nemacol-1 and Nemacol-5) to have low micromolar activity against *Dirofilaria* (Fig. 3). Of the 20 analogs tested that contained a 4-nitrophenyl group, 10 were active against *D. immitis* (LC50 ≤ 22.5 μM). Furthermore, only analogs containing 4-nitrophenyl in position R_1_ were active against *D. immitis*. Together, these observations suggest that the nitrophenyl group is important for activity against *D. immitis*. These results show that the Nemacol scaffold has activity beyond *C. elegans*.

Next, we chose 12 diversely structured analogs that had good activity against *C. elegans* and investigated whether any might have weakened affinity for the rat VAChT relative to vesamicol and Nemacol-1. Ten of the 12 analogs tested had more than 10-fold less affinity to the rat VAChT relative to vesamicol and five had more than 30-fold less affinity (Fig. 3b and 3c).

Comparing Nemacol-1’s *in vitro* mammalian VAChT Ki (0.66 µM) to its *C. elegans in vivo* EC50 activity (7.0 µM) reveals an activity ratio of 0.09. By contrast, Nemacol-63’s equivalent ratio is 1.28, which is an improvement of over 14-fold in potential selectivity. The two analogs with the poorest VAChT affinity (Nemacol-62 and 63) share a cyclopentylmethyl group in the R_2_ position, suggesting that this feature diminishes affinity with mammalian VAChT. Together, the results suggest that it may be possible to modify the Nemacol scaffold to achieve nematode selectivity.

### Nemacol Synthetically Interacts with Ivermectin

The nAChR antagonist and anthelmintic derquantel synthetically interacts with abamectin, a macrocyclic lactone and Cl^-^ channel agonist^22,23,39-41^. Nemacol inhibits VAChT and consequently depresses cholinergic signaling by decreasing synaptic vesicle ACh content. We reasoned that Nemacol, like derquantel, might therefore synthetically interact with macrocyclic lactones to disrupt nematode neuromuscular function. Indeed, we found that in three-day liquid development assays (see methods), combinations of ivermectin and Nemacol could yield effective killing of *C. elegans* at concentrations that had little effect on their own (Fig. 4a,b). The combination yields a global <u>Z</u>ero Interaction <u>P</u>otency (ZIP) synergy score of 20.0, which is beyond the ZIP score threshold of synergy (10)^42-44^ (Fig. 4c). Ivermectin is a known and potent drug pump inhibitor and may block the worm’s elimination of Nemacol^45,46^.

**Figure 4.**
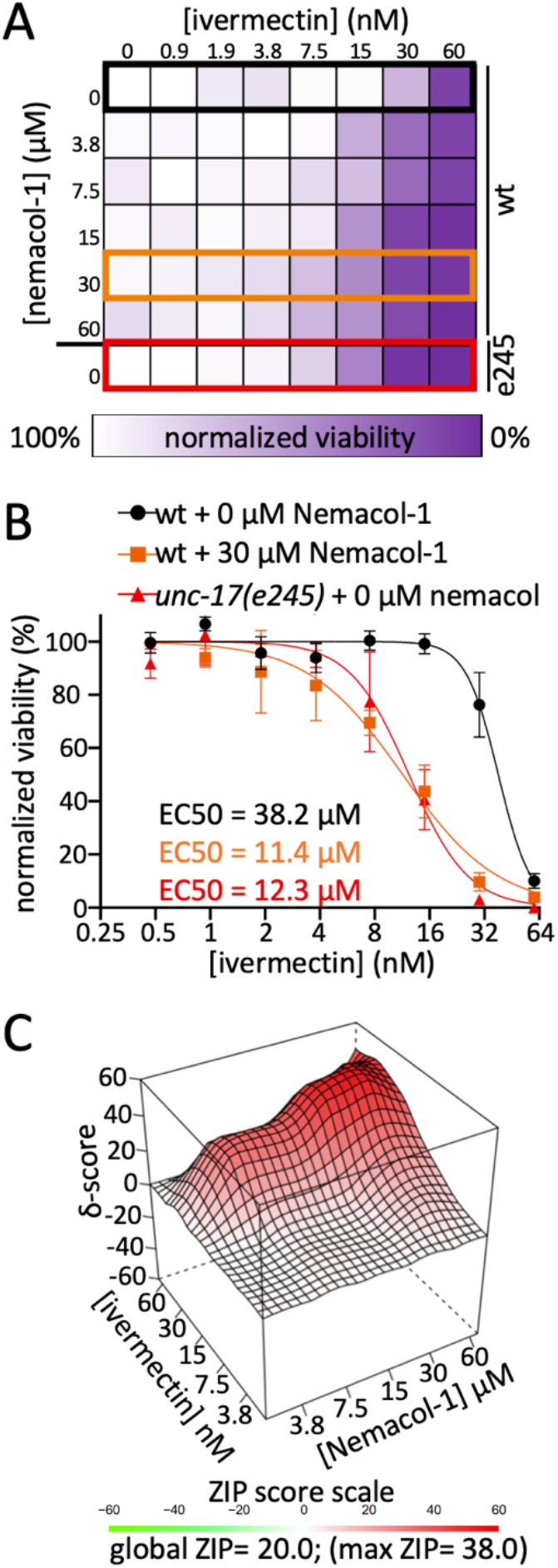
Nemacol-1 synergistically kills *C. elegans* in combination with ivermectin. (**A**) *C. elegans* viability over a double dose response matrix of Nemacol-1 and ivermectin reporting the control normalized fraction of animals alive in treatment wells of 3-days of growth assays (see methods). ∼20 *C. elegans* L1s were added to wells containing the indicated condition with 0.6% DMSO and an HB101 *E. coli* food source in nematode growth medium. Data are the calculated mean from three biological replicates conducted in technical duplicate. (**B**) Dose-response curves from panel A highlighting the shift of ivermectin *C. elegans* killing +/- 30 μM Nemacol-1 in wild-type (black versus orange), or wild-type versus *unc-17(e245)* mutant animals (black versus red). For wt + 0 μM Nemacol-1 compared to wt + 30 μM Nemacol-1 *p* = 1.0 × 10^−15^; for wt + 0 μM Nemacol-1 compared to *unc-17(e245)* + 0 μM Nemacol-1 *p =* 5.0 × 10^−15^. *P*-value calculated using an extra sum-of-squares F test comparing the wild-type ivermectin dose response to the indicated dose response in GraphPad Prism version 9.3.1. (**C**) Zero Interaction Potency (ZIP) synergy score plot of the ivermectin + Nemacol-1 double-dose response interaction generated using the SynergyFinder2.0 server ^44^. The combination yields global ZIP synergy score of 20.0, which is beyond the ZIP score threshold of synergy (10)^33-35^.

To test whether the inhibition of VAChT contributed to the synthetic interaction between Nemacol and ivermectin, we tested whether the *unc-17(e245)* reduction-of-function mutant was more sensitive to the effects of ivermectin (Fig.4a). Indeed, *unc-17(e245)* demonstrated a sensitivity to ivermectin that was comparable to 30 μM Nemacol (Fig. 4b). This suggests that the synergistic interaction between Nemacol and ivermectin is not solely due to altered metabolism or elimination induced by ivermectin. These data indicate that Nemacol may have additional utility in its ability to sensitize nematodes to one of the most widely used anthelmintics in the world.

## Discussion

The current repertoire of anthelmintics available for the control of parasitic nematodes that infect humans and non-human animals is lacking^1,47^. Hence, the identification of novel and selective nematicidal compounds is a key step in protecting human health and food security. Here, we have identified the Nemacol scaffold that disrupts nematode motor function via the inhibition of VAChT. Select Nemacol analogs maintain nematode activity whilst losing affinity for mammalian VAChT, suggesting that nematode selectivity can be achieved with this scaffold. Nemacol is detoxified in *C. elegans* over the course of 24 hours. However, assays performed against multiple parasitic nematodes over an equivalent timespan (or longer) indicate that Nemacol’s effects can perdure in parasites.

How does Nemacol compare to other VAChT inhibitors in the context of being a candidate anthelmintic lead? Nemacol is similar in structure to vesamicol, a canonical VAChT inhibitor^48^ and both compounds likely interact with a common binding site on VAChT^32^. However, the vesamicol scaffold has lackluster activity against nematodes, has high affinity for mammalian VAChT, and has low tolerability in rats^27,32,49^. These features likely account for a lack of interest in developing the vesamicol scaffold as an anthelmintic.

In contrast to vesamicol, the spiroindoline scaffold has been rigorously pursued as a candidate anthelmintic because it selectively incapacitates nematodes and insects^28,31,50^. Indeed, there has been heavy commercial interest in pursuing the spiroindolines as candidate anthelmintics by several groups, including Zoetis Services LLC^28^, Syngenta Ltd^30^ and Intervet Inc^29^. The spiroindolines have been shown to inhibit VAChT via residues that are at least partially distinct from those that interact with vesamicol^31,32^. Of note, Nemacol analogs incapacitate *C. elegans* motor activity at equivalent concentrations to the spiroindolines (compare Figure 6 in Sluder et al., 2012 to Fig. 3a; note 1 μg/mL SYN351 = 1.9 μM) and Nemacol analogs have also proved comparatively effective at incapacitating *D. immitis* filariae relative to spiroindolines (compare Table 2 in US20160296499A1 to Fig. 3a; note we report LD50 compared to minimal inhibitory concentration^28^).

The high lipophilicity of the spiroindolines may have so far stifled their development into commercial products. Guidance from the European Union’s European Chemical Agency^51^ highlights that compounds with a Log P greater than 4 have accumulative potential in adipose tissue of animals. The primary spiroindoline lead pursued by Syngenta (SYN876 in Sluder et al., 2012^27^) has a SwissADME predicted consensus LogP of 5.53 suggesting that this lead may have concerning accumulative potential^52^. In contrast, 90% of Nemacol analogs had a SwissADME predicted consensus Log P < 4.0 with a median of 2.88^52^.

Relative to the structural complexity of the spiroindoline scaffold (MW 453.53, synthetic accessibility score of 4.23)^52^ the Nemacol-1 structure is simpler (MW 303.4) with a synthetic accessibility score of 2.52^52,53^). The higher the synthetic accessibility score, the more difficult the synthesis^52,53^. Indeed, we have found Nemacol to have a relatively inexpensive synthesis route with several analogs so far synthesized requiring only a two-step metal-free synthetic sequence that doesn’t require purification of the intermediate (see methods).

VAChT has clearly been recognized by multiple commercial groups as an attractive anthelmintic target^27-30^. One reason for this is it may be difficult to mutate VAChT to a state that reduces an inhibitor’s efficacy without compromising the transporter itself. VAChT mutant residues that confer resistance to vesamicol or the spiroindolines disrupt the ability of these molecules to interact with VAChT, but reduce acetylcholine interaction and lead to obvious locomotory defects in *C. elegans*^27,32,54,55^. Missense mutations that alter the ability of VAChT inhibitors to interact with the transporter will likely confer a clear selective disadvantage to worms. A second reason for the keen interest in the VAChT target may be because of its high conservation across nematodes^56^ (Fig. S3). Despite the conservation, there are key differences in nematode VAChT sequence relative to non-target species near the presumptive vesamicol binding site, suggesting that broad-spectrum nematode-selectivity may be possible to achieve with VAChT inhibitors (Fig. S3). Given Nemacol’s synthetic accessibility and the attractiveness of its target, Nemacol represents an important scaffold to further develop as a novel anthelmintic molecule.

## Methods

### Worm Culture and Strains

All nematode strains were cultured using standard methods at 20°C unless otherwise indicated^57^. The N2 (wild-type) strain of *Caenorhabditis elegans* and *C. elegans* mutant strains were obtained from the *C. elegans* Genetic Center (University of Minnesota). Synchronization of worm developmental stages was achieved using standard bleach preparation protocols^58^. Worms were maintained on Modified Youngren’s, Only Bacto-peptone (MYOB) media containing 2% agar with a surface lawn of OP50 strain *Escherichia coli*^59^.

### Preparation of small-molecules in solid media

Molten Modified Youngren’s, Only Bacto-peptone (MYOB) containing 2% agar was equilibrated to 55°C in a water bath. Small-molecules solvated in DMSO were spiked into at least 4 mL of molten media mixture in 15 mL conical tubes, inverted 5 times and vortexed. The final concentration of dimethyl sulfoxide (DMSO) in each of the wells was 1% v/v. One mL of media containing small-molecule was pipetted into wells of a 24-well plate using a 10 mL Sarstedt serological pipette. Plates were dried under sterile air flow in a laminar flow hood for 90 minutes. After 90 minutes 25 μL of OP50 strain *E. coli* bacteria culture from a saturated Luria Broth culture was added by pipette onto the surface of the culture media. Plates were allowed to dry on a benchtop proximal to flame for 15 minutes. Plates were covered and wrapped in tinfoil and were used the following day.

### Scoring of Motor Phenotypes

Locomotor phenotype analyses were done in 24-well plates with 1 mL of MYOB substrate (27.5 g Trizma HCl, 12 g Trizma Base, 230 g bacto tryptone, 10 g NaCl, 0.4 g cholesterol (95%)) seeded with 25 µL of OP50 *Escherichia coli* on each well. Each compound was added to the MYOB substrate before pouring to achieve the desired final concentrations of 30 µM or 60 µM after diffusion through the media. Young adult worms are transferred into each well using a platinum wire pick. A Leica MZ75 stereomicroscope was used to visualize the movement of worms on the solid substrate. The specific dominant locomotor phenotype (i.e. ‘jerky-uncoordinated’, ‘paralyzed’, ‘coiler’ or ‘wild-type locomotion’) was scored after the touch on the head with a platinum wire over a ∼3-5 seconds of observation. Animals were scored as jerky-Unc if animals exhibit a lack of smooth locomotion with frequent abrupt stops and start before and after a touch on the head; animals were scored as coiler if animals are observed in a coiled position (Supplemental Movie 1); animals were scored as paralyzed if animals exhibit flaccid paralysis and fail to reverse upon light touch of the head with a platinum wire; animals were scored as ‘wild-type locomotion’ if animals exhibit normal sinusoidal locomotion and/or a normal backing response upon light touch on the head with a platinum wire.

### C. elegans 3-day development assays

*C. elegans* larval development assays were conducted in 96-well flatbottom clear flatbottom plates. ∼20 L1 larvae in 10 μL of M9 buffer were pipetted into each test wells containing 40 μL of Nematode Growth Media (NGM)^60^ media supplemented with HB101 *E. coli* with the desired test compound (+0.6% dimethylsulfoxide (DMSO) as the chemical solvent). Plates were wrapped in 3 layers of brown paper towels soaked with water. After either 3 days of incubation the number of *C. elegans* animals of different larval stages were recorded using a Leica MZ75 stereomicroscope.

### Chemistry

#### General Considerations

Unless otherwise stated, all reactions were carried out under an atmosphere of dry argon, using glassware that was either oven (120 °C) or flame-dried. Work-up and isolation of compounds was performed using standard benchtop techniques. All commercial reagents were purchased from chemical suppliers (Sigma-Aldrich, Combi-Blocks, or Alfa Aesar) and used without further purification. Dry solvents were obtained using standard procedures (dichloromethane and acetonitrile were distilled over calcium hydride). Reactions were monitored using thin-layer chromatography (TLC) on EMD Silica Gel 60 F254 plates. Visualization was performed under UV light (254 nm) or using potassium permanganate (KMnO_4_) stain. Flash column chromatography was performed on Siliaflash P60 40-63 µm silica gel purchased from Silicycle. NMR characterization data was obtained at 293 K on a Varian Mercury 300 MHz, Varian Mercury 400 MHz, Bruker Advance III 400 MHz, Agilent DD2 500 MHz equipped with a 5 mm Xses cold probe or Agilent DD2 600 MHz. ^1^H spectra were referenced to the residual solvent signal (CDCl_3_ = 7.26 ppm, DMSO-*d*_*6*_ = 2.50 ppm). ^13^C{^1^H} spectra were referenced to the residual solvent signal (CDCl_3_ = 77.16 ppm, DMSO-*d*_*6*_ = 39.52 ppm). Data for ^1^H NMR are reported as follows: chemical shift (δ ppm), multiplicity (s = singlet, d = doublet, t = triplet, q = quartet, m = multiplet), coupling constant (Hz), integration. NMR spectra were recorded at the University of Toronto

Department of Chemistry NMR facility (http://www.chem.utoronto.ca/facilities/nmr/nmr.html). Infrared spectra were recorded on a Perkin-Elmer Spectrum 100 instrument equipped with a single-bounce diamond/ZnSe ATR accessory in the solid state and are reported in wavenumber (cm^-1^) units. High resolution mass spectra (HRMS) were recorded at the Advanced Instrumentation for Molecular Structure (AIMS) in the Department of Chemistry at the University of Toronto (https://www.chem.utoronto.ca/chemistry/AIMS.php).

#### General Procedures

##### General Procedure A

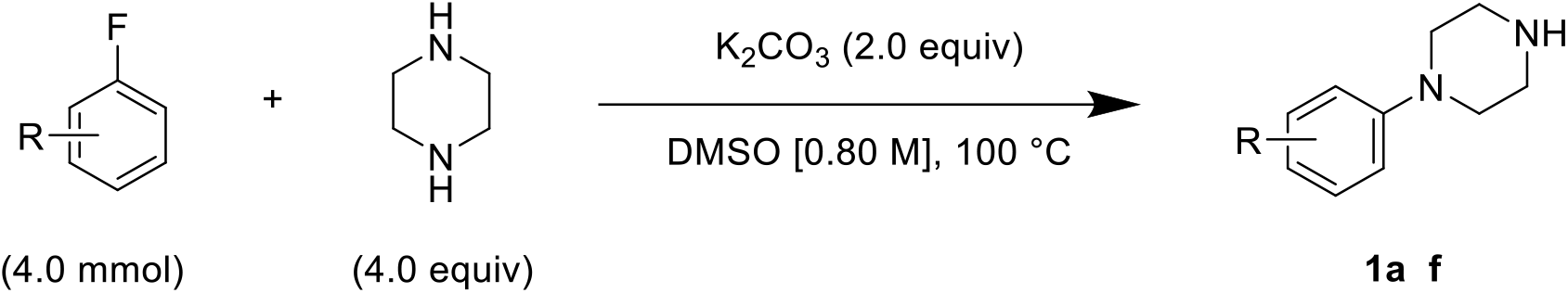

The procedure was modified from literature ^61^. To a round-bottom flask at room temperature were added piperazine (1.4 g, 16 mmol, 4.0 equiv), potassium carbonate (1.1 g, 8.0 mmol, 2.0 equiv), dimethyl sulfoxide (4.7 mL) then the corresponding fluorobenzene derivative (4.0 mmol, 1.0 equiv) and the mixture was stirred at 100 °C. Once the reaction was complete as indicated by TLC (approximately 22 hours), the reaction mixture was diluted with ethyl acetate, transferred to a separatory funnel and washed with water (3 times) then saturated aqueous sodium chloride. The organic phase was dried with Na_2_SO_4_, filtered, then concentrated on a rotary evaporator and the substituted 1-phenylpiperazine (**1a–f**) was obtained. The crude solid was used in the next step without further purification.

##### General Procedure B

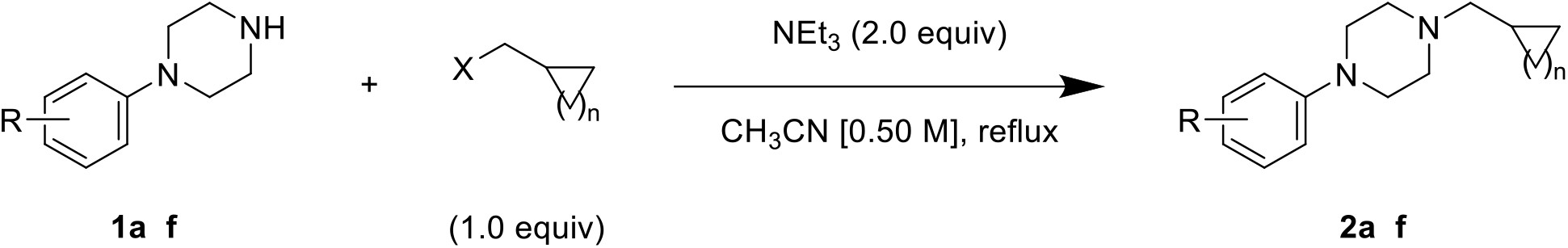

To a round-bottom flask were added **1** (0.50 mmol, 1.0 equiv), dry acetonitrile (1.0 mL), triethylamine (0.14 mL, 1.0 mmol, 2.0 equiv) then the alkyl halide or toluenesulfonate (0.50 mmol, 1.0 equiv) and the mixture was stirred at reflux until completion as indicated by TLC (approximately 24 hours). The mixture was diluted with ethyl acetate, transferred to a separatory funnel and washed with water (3 times) then saturated aqueous sodium chloride. The organic phases were combined and dried with Na_2_SO_4_, filtered, and concentrated on a rotary evaporator. The residue was purified by column chromatography with the indicated eluent and the alkylated arylpiperazine (**2**) was obtained.

##### General Procedure C

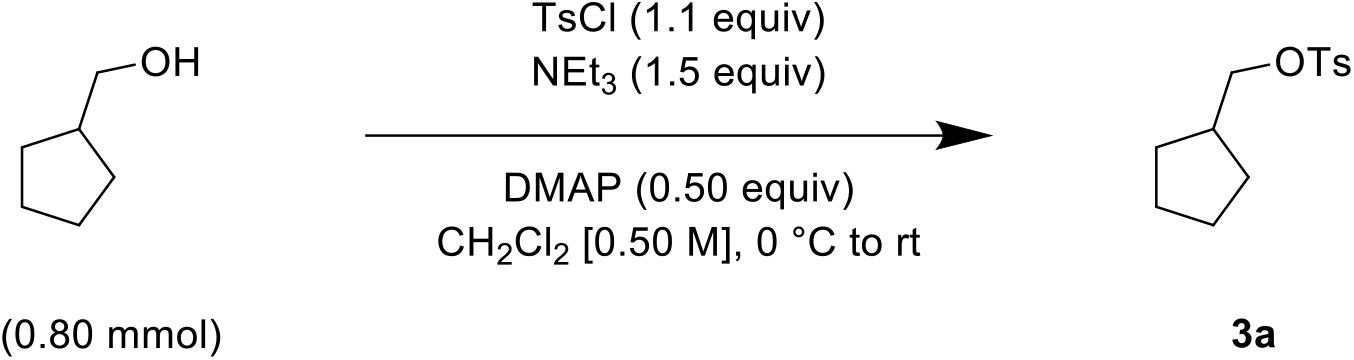

The procedure was modified from literature^61^. To a round-bottom flask were added cyclopentylmethanol (0.087 mL, 0.80 mmol, 1.0 equiv) and dry dichloromethane (1.6 mL) and the flask was submerged in an ice-water bath. Triethylamine (0.17 mL, 1.2 mmol, 1.5 equiv) was added to the flask followed by 4-(dimethylamino)pyridine (48.9 mg, 0.40 mmol, 0.50 equiv) then *p*-toluenesulfonyl chloride (168 mg, 0.88 mmol, 1.1 equiv) and the mixture was warmed to room temperature and stirred until completion as indicated by TLC (approximately 3 hours). Dichloromethane and water were added, then the mixture was transferred to a separatory funnel and the layers were separated. The organic phase was washed with water (2 times) followed by saturated aqueous sodium chloride, then dried over Na_2_SO_4_, filtered, and concentrated on a rotary evaporator. The residue was purified by column chromatography (19/1 v/v pentanes/ethyl acetate) and cyclopentylmethyl 4-methylbenzenesulfonate (**3a**, 59% yield) was obtained as a colourless oil. The spectral data were in accordance with literature.

#### Characterization Data for Products

##### 4-(4-(cyclohexylmethyl)piperazin-1-yl)benzonitrile (2a, Nemacol-22)

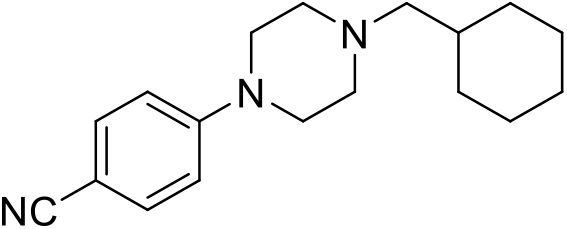

Synthesized according to General Procedures A and B from 4-fluorobenzonitrile and (bromomethyl)cyclohexane. Compound **2a** was isolated as a bright yellow solid (MP = 61–63 °C). ^**1**^**H NMR (CDCl**_**3**_, **500 MHz):** δ 7.46 (d, *J* = 9.0 Hz, 2H), 6.83 (d, *J* = 9.0 Hz, 2H), 3.32 – 3.28 (m, 4H), 2.52 – 2.48 (m, 4H), 2.15 (d, *J* = 7.2 Hz, 2H), 1.78 (dddd, *J* = 14.1, 4.9, 3.5, 1.5 Hz, 2H), 1.74 – 1.63 (m, 3H), 1.50 (ttt, *J* = 10.8, 7.2, 3.5 Hz, 1H), 1.28 – 1.11 (m, 3H), 0.93 – 0.83 (m, 2H). ^**13**^**C NMR (CDCl**_**3**_, **125 MHz):** δ 153.6, 133.5, 120.2, 114.1, 99.9, 65.6, 53.3, 47.2, 35.1, 31.9, 26.9, 26.2. **IR (neat):** 2922, 2849, 2780, 2216, 1599, 1515, 1447, 1247, 1177, 818. **HRMS (DART):** calc. for C_18_H_26_N_3_ 284.2127 [M+H]^+^, found 284.2132.

##### 1-(cyclohexylmethyl)-4-(2-methyl-4-nitrophenyl)piperazine (2b, Nemacol-23)

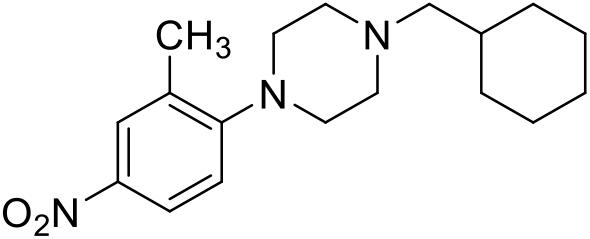

Synthesized according to General Procedures A and B from 1-fluoro-2-methyl-4-nitrobenzene and (bromomethyl)cyclohexane. Compound **2b** was isolated as a bright yellow solid (MP = 52–54 °C). ^**1**^**H NMR (CDCl**_**3**_, **400 MHz):** δ 8.05 – 8.00 (m, 2H), 6.98 (dt, *J* = 8.5, 1.1 Hz, 1H), 3.04 (t, *J* = 4.8 Hz, 4H), 2.61 – 2.52 (m, 4H), 2.35 (s, 3H), 2.20 (d, *J* = 7.1 Hz, 2H), 1.85 – 1.64 (m, 5H), 1.59 – 1.47 (m, 1H), 1.32 – 1.14 (m, 3H), 0.90 (q, *J* = 12.5 Hz, 2H). ^**13**^**C NMR (CDCl**_**3**_, **125 MHz):** δ 157.7, 142.3, 132.2, 126.8, 122.8, 118.2, 65.7, 53.8, 51.2, 35.1, 32.0, 26.9, 26.3, 19.0. **IR (neat):** 2918, 2842, 2810, 1583, 1504, 1447, 1328, 1228, 1011, 893. **HRMS (DART):** calc. for C_18_H_28_N_3_O_2_ 318.2182 [M+H]^+^, found 318.2178.

##### 1-(cyclohexylmethyl)-4-(3-methyl-4-nitrophenyl)piperazine (2c, Nemacol-24)

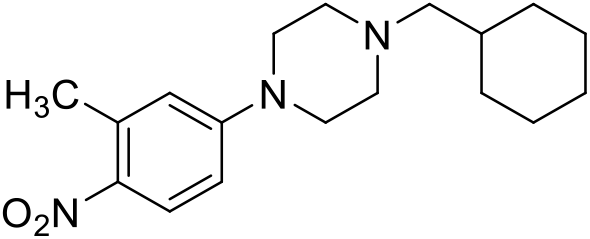

Synthesized according to General Procedures A and B from 4-fluoro-2-methyl-1-nitrobenzene and (bromomethyl)cyclohexane. Compound **2c** was isolated as a bright yellow solid (MP = 75–77 °C). ^**1**^**H NMR (CDCl**_**3**_, **500 MHz):** δ 8.06 (d, *J* = 9.3 Hz, 1H), 6.68 (dd, *J* = 9.3, 2.9 Hz, 1H), 6.61 (d, *J* = 2.9 Hz, 1H), 3.39 – 3.35 (m, 4H), 2.62 (s, 3H), 2.53 – 2.49 (m, 4H), 2.16 (d, *J* = 7.2 Hz, 2H), 1.82 – 1.75 (m, 2H), 1.75 – 1.64 (m, 3H), 1.50 (ttt, *J* = 10.8, 7.2, 3.5 Hz, 1H), 1.29 – 1.12 (m, 3H), 0.94 – 0.83 (m, 2H). ^**13**^**C NMR (CDCl**_**3**_, **125 MHz):** δ 154.0, 139.0, 137.3, 127.9, 116.2, 111.2, 65.6, 53.3, 47.1, 35.1, 32.0, 26.9, 26.2, 22.8. **IR (neat):** 2920, 2843, 1638, 1600, 1533, 1249, 1226, 1001, 925, 822. **HRMS (DART):** calc. for C_18_H_28_N_3_O_2_ 318.2182 [M+H]^+^, found 318.2189.

##### 1-(cyclohexylmethyl)-4-(pyridin-4-yl)piperazine (2d, Nemacol-25)

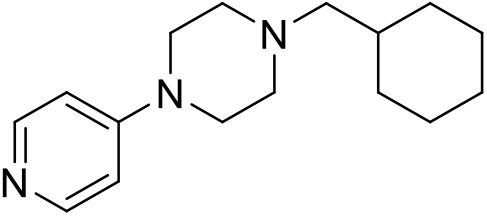

Synthesized according to General Procedures A and B from 4-chloropyridine hydrochloride and (bromomethyl)cyclohexane. Compound **2d** was isolated as a white solid (MP = 66–70 °C). ^**1**^**H NMR (DMSO-*d***_***6***_, **500 MHz):** δ 8.17 (d, *J* = 6.1 Hz, 2H), 6.92 (d, *J* = 6.1 Hz, 2H), 3.40 (t, *J* = 5.1 Hz, 4H), 2.43 (t, *J* = 5.1 Hz, 4H), 2.12 (d, *J* = 7.2 Hz, 2H), 1.76 – 1.70 (m, 2H), 1.69 – 1.58 (m, 3H), 1.50 (ttt, *J* = 10.9, 7.3, 3.4 Hz, 1H), 1.26 – 1.07 (m, 3H), 0.89 – 0.77 (m, 2H). ^**13**^**C NMR (DMSO-*d***_***6***_, **125 MHz):** δ 155.2, 146.3, 108.0, 64.6, 52.6, 45.5, 34.2, 31.2, 26.4, 25.5. **IR (neat):** 2919, 2849, 1638, 1600, 1533, 1249, 1132, 1001, 989, 805. **HRMS (DART):** calc. for C_16_H_26_N_3_ 260.21267 [M+H]^+^, found 260.2120.

##### 4-(4-(cyclopentylmethyl)piperazin-1-yl)benzonitrile (2e, Nemacol-27)

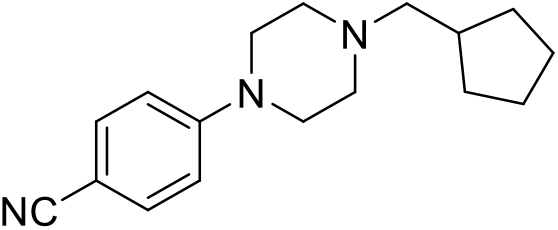

Synthesized according to General Procedures A, B and C from 4-fluorobenzonitrile and **3a**. Compound **2e** was isolated as a light orange solid (MP = 49–51 °C). ^**1**^**H NMR (CDCl**_**3**_, **500 MHz):** δ 7.47 (d, *J* = 9.0 Hz, 2H), 6.84 (d, *J* = 9.0 Hz, 2H), 3.34 – 3.29 (m, 4H), 2.59 – 2.52 (m, 4H), 2.30 (d, *J* = 7.3 Hz, 2H), 2.08 (hept, *J* = 7.3 Hz, 1H), 1.80 – 1.71 (m, 2H), 1.65 – 1.48 (m, 4H), 1.26 – 1.16 (m, 2H). ^**13**^**C NMR (CDCl**_**3**_, **125 MHz):** δ 153.6, 133.6, 120.3, 114.1, 100.0, 64.5, 53.1, 47.2, 37.2, 31.5, 25.3. **IR (neat):** 2947, 2936, 2864, 2816, 2211, 1601, 1513, 1448,1166, 921. **HRMS (DART):** calc. for C_17_H_24_N_3_ 270.19702 [M+H]^+^, found 270.1968.

##### 1-(cyclopropylmethyl)-4-(4-nitrophenyl)piperazine (2f, Nemacol-28)

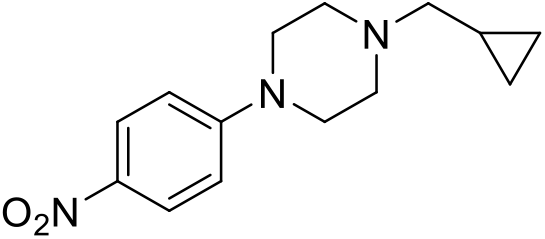

Synthesized according to General Procedures A and B from 1-fluoro-4-nitrobenzene and (bromomethyl)cyclopropane. Compound **2f** was isolated as a dark orange solid. The characterization data are in accordance with literature^62^. ^**1**^**H NMR (CDCl**_**3**_, **500 MHz):** δ 8.12 (d, *J* = 9.4 Hz, 2H), 6.82 (d, *J* = 9.4 Hz, 2H), 3.46 (t, *J* = 5.2 Hz, 4H), 2.68 (t, *J* = 5.2 Hz, 4H), 2.32 (d, *J* = 6.6 Hz, 2H), 0.90 (ttt, *J* = 8.0, 6.6, 4.9 Hz, 1H), 0.58 – 0.54 (m, 2H), 0.16 – 0.12 (m, 2H). ^**13**^**C NMR (CDCl**_**3**_, **125 MHz):** δ 155.0, 138.5, 126.1, 112.7, 63.7, 52.8, 47.1, 8.4, 4.1.

### Commercial Sources of Nemacol Phenylpiperazines

Nemacol analogs were procured from several commercial sources as dry compound. Molecules were sourced from Chembridge Inc., Enamine and OTAVA chemicals, Ltd. Vesamicol was purchased from SigmaAldrich as (±)-Vesamicol hydrochloride (product ID V100).

### Radioligand Competition Assays for Evaluation of Phenylpiperazine VAChT Affinity

Rat VAChT K_i_ data are the K_i_ of nemacol analog as determined from radioligand [^3^H]vesamicol binding assays using PC12 cells expressing ratVAChT. [^3^H]vesamicol was procured from PerkinElmer (product ID: AH5183 (L-[Piperidinyl-3,4-3H]-(Vesamicol), 250 µCi)). Stably transfected PC12 cells expressing ratVAChT were shared by Dr. Ali Roghani and radioligand competition experiments conducted using standard protocols^63,64^.

### Parasitic Nematode Culture and Small-Molecule Assays

*Dirofilaria immitis* microfilariae and larval stage 3 (L3) worms were performed in the laboratories of Bayer Animal Health GmbH (Monheim, Germany) in accordance with the local Animal Care and Use Committee and governmental authorities (LANUV#200/A176 and #200/A154). For microfilariae immobility assays, approximately 250 freshly purified microfilariae were cultured in single wells of a 96-well microtiter plate containing supplemented RPMI 1640 medium. Compounds were added in the following concentrations:50 µM, 10 µM, 2 µM, 0.4 µM, 0.08 µM, 0.016 µM and 0.0032 µM. Microfilariae exposed to medium substituted with 1% DMSO were used as negative controls. Motility of microfilariae was evaluated after 72 hours of drug exposure using an image-based approach – DiroImager, developed by Bayer Technology Services. Data are reported as the EC50 (µM) calculated from the tested concentration series. For L3 development assays, freshly isolatedL3s were cultured in wells of a 96-well microtiter plate with 10 L3 per well. All wells contained supplemented RPMI 1640 medium and test compound at one of the following concentrations: 10 µM, 2 µM, 0.4 µM, 0.08 µM, 0.016 µM and 0.0032 µM. L3s exposed to DMSO only (1%) were used as negative controls. All drug concentrations were tested in duplicate and drug effects were evaluated after 72 hours of incubation. Motility was scored for each individual worm. Each motile worm, independently of the degree of motility, reduced the drug activity by 5%, as two cavities with 10 worms each (20 worms in total = 100% activity if all of them are completely paralyzed) were tested per indicated compound. Data were only considered as valid if at least 90% worms in the negative control group remained as motile as observed at the beginning of the experiment. Data are reported as the EC50 (µM) calculated from the tested concentration series.

### Statistical Analyses

Unpaired one or two sided t-tests, or Chi-square tests were conducted between control and treatment groups where appropriate and as indicated in figure legends. Extra sum-of-squares F tests were conducted comparing EC50 curves generated for dose-response data in GraphPad Prism (version 9.3.1).

### emb-8 RNAi High Performance Liquid Chromatography Coupled to a Diode Array Detector (HPLC-DAD) analyses

Young adult worms, grown from synchronized first-staged larval hatchlings at 25 °C for 48 hours were used for the HPLC accumulation analyses. The worms were grown on either OP50 *E. coli*, or HT115 *E. coli* expressing RNAi plasmid for *emb-8* knockdown. 2000 worms were plated on MYOB agar plates containing the chemical of interest (1% DMSO v/v). RNAi by feeding assays were performed on MYOB agar plates seeded with bacteria expressing the *emb-8* RNAi. The cultivation and induction followed the principles outlined by Hull and Timmons^65^. The strains used were the HT115 *E. coli* feeding strain, with the *emb-8* dsRNA cloned into the L4440 plasmid, and a control strain with the empty L4440 plasmid. Liquid bacterial cultures were cultivated overnight in the presence of 100 µg/mL ampicillin to saturation. The following day, 6 cm MYOB agar plates with final concentrations of 100 µg/mL carbenicilin and 1 mM IPTG were seeded with 250 µL of the bacterial culture and dried at room temperature overnight. First larval-staged wildtype and emb*-8*[*(hc69)ts*] worms which were grown and hatched at 15 °C, were plated on the MYOB RNAi plates. These worms were grown at 25 °C and allowed to feed for 2 days before being transferred to experimental plates. All experimental chemical plates contained final concentrations of 100 µg/mL carbenicilin and 1 mM IPTG and were seeded with *emb-8* RNAi and L4440 control bacteria. Worms were incubated on drug plates for 6 hours at 25 °C unless otherwise noted. After incubation, worms were washed off the agar plates and suspended in M9 buffer containing 0.5% gelatin^66^. After three washes in gelatin-M9, 500 µL of worm suspension was added to each well of Pall ACropPrep 96-well filter plates (0.45 µm GHP membrane, 1 mL well volume). The buffer was drained from the wells by vacuum and the worms were resuspended in 50 µL of M9 buffer and frozen at -80 C. The samples were later lysed by adding 35 μl of a 2X lysis solution (100 mM KCl, 20 mM Tris, pH 8.3, 0.4% SDS, 120 μg ml-1 proteinase K) to each tube at 56 °C for 1 h. Prior to HPLC coupled to a diode array detector (HPLC-DAD) analyses analysis, 70µL of acetonitrile was added to the lysates. The samples were then mixed by vortexing for 10 seconds, and centrifuged at 17,949 *g* for 2 minutes. After centrifugation, 100 µL of the lysate was injected onto a 4.6 × 150 mm Zorbax SB-38 column (5 μm particle size) and eluted with solvent and flow rate gradients over 5.2 minutes. UV-Vis absorbance was measured every 2 nm between 190 and 602 nm. Using MATLAB (The MathWorks), absorbance intensity values were converted to three-dimensional heat-mapped chromatograms. A sample of 5 nmol pure Nemacol-1 was processed prior to worm samples to determine the compounds elution time and absorbance spectrum. All HPLC was performed a using an HP 1050 system equipped with an autosampler, vacuum degasser, and a variable wavelength diode-array detector. The solvent and flow rate gradients are indicated below in Table S1. Data analysis was done using HP Chemstation software. The area under the curve was automatically integrated by the software for quantification of Nemacol-1 absorbance peaks. For each sample, the ratio of parent and individual compound metabolite (M1, M2 and M3) area under the curve to the total area under the curve for all compound related peaks (parent, M1, M2 and M3) were calculated. As an example, for the calculation of the parent compound ratio, the area under the curve for the parent compound was divided by the total area under the curve for the parent compound and all metabolite related peaks.

---

**Table S1:**
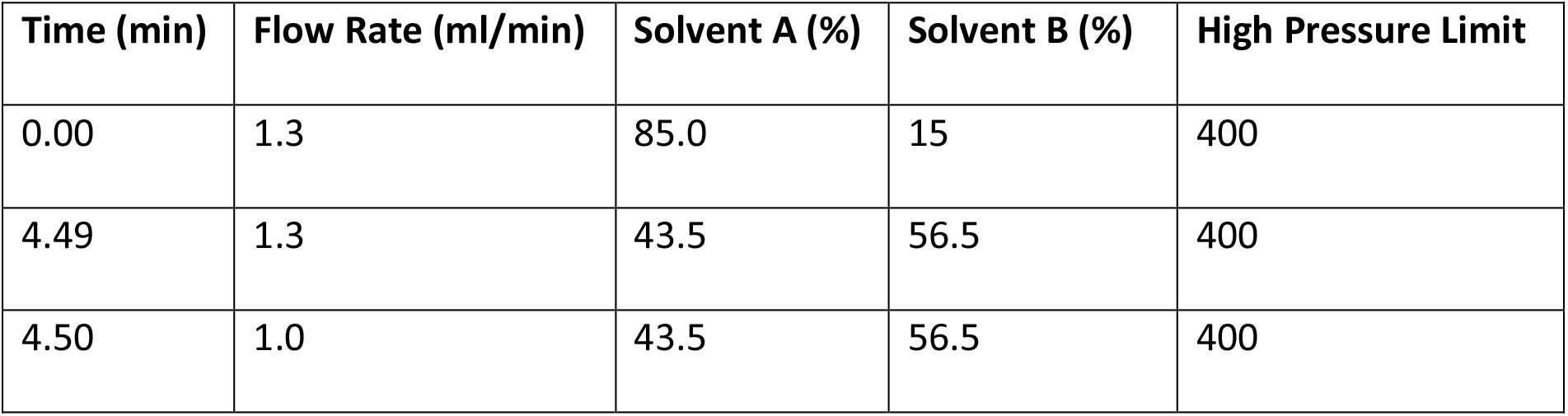
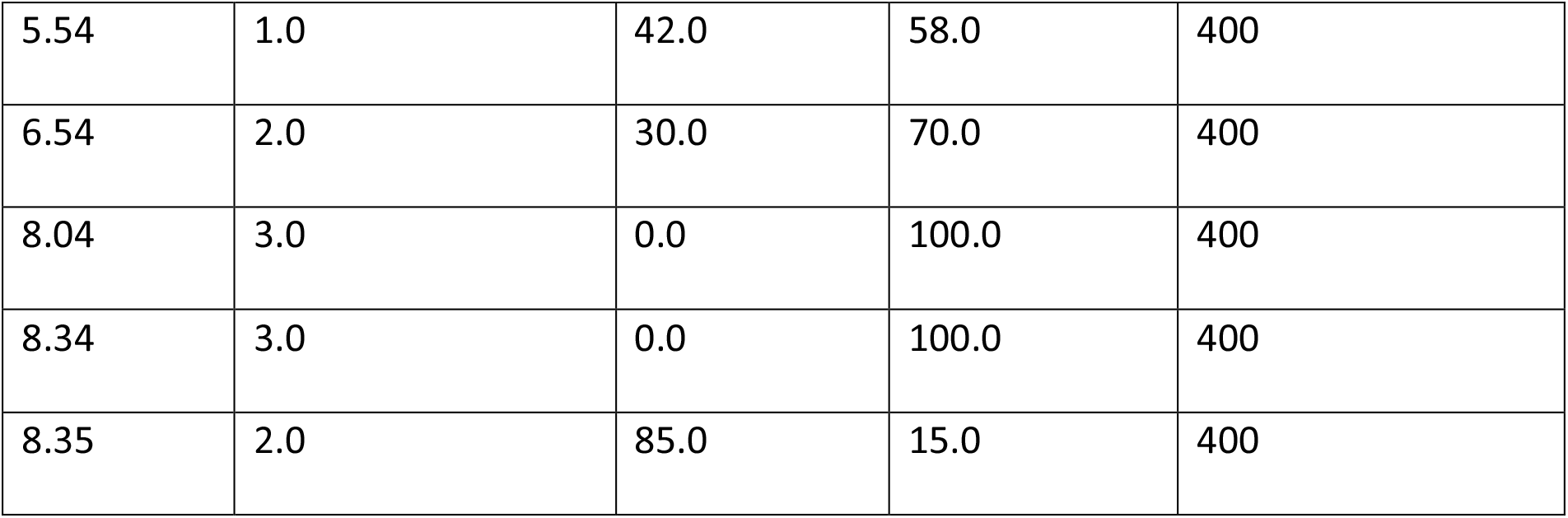
HPLC solvent and flow rate gradients.

## Acknowledgements

We are grateful for the strains given to us by the *Caenorhabditis* Genetics Center (University of Minnesota) and James Rand Oklahoma (Center for Neuroscience).

## Ethics Statement

DK: Experiments on *D. immitis* were performed in the laboratories of Bayer Animal Health GmbH (Monheim, Germany) in accordance with the local Animal Care and Use Committee and governmental authorities.

## Author Contributions

Conceptualization: PJR, SH, JP, ARB

Methodology: SH, JP, TS, RJB, ML, DK, WD-C

Investigation: SH, JP, TS, RJB, JC, DK, WD-C

Visualization: PJR, SH, JP

Funding acquisition: PJR

Project administration: PJR

Supervision: PJR, ML, DK, WD-C, PB

Writing – original draft: SH, JP, PJR

Writing – review & editing: PJR, SH, JP, RJB, ML, PB

## Data and materials availability

Original data for all analyses presented are included in the Supplementary Data File.

---

**Figure S1.**
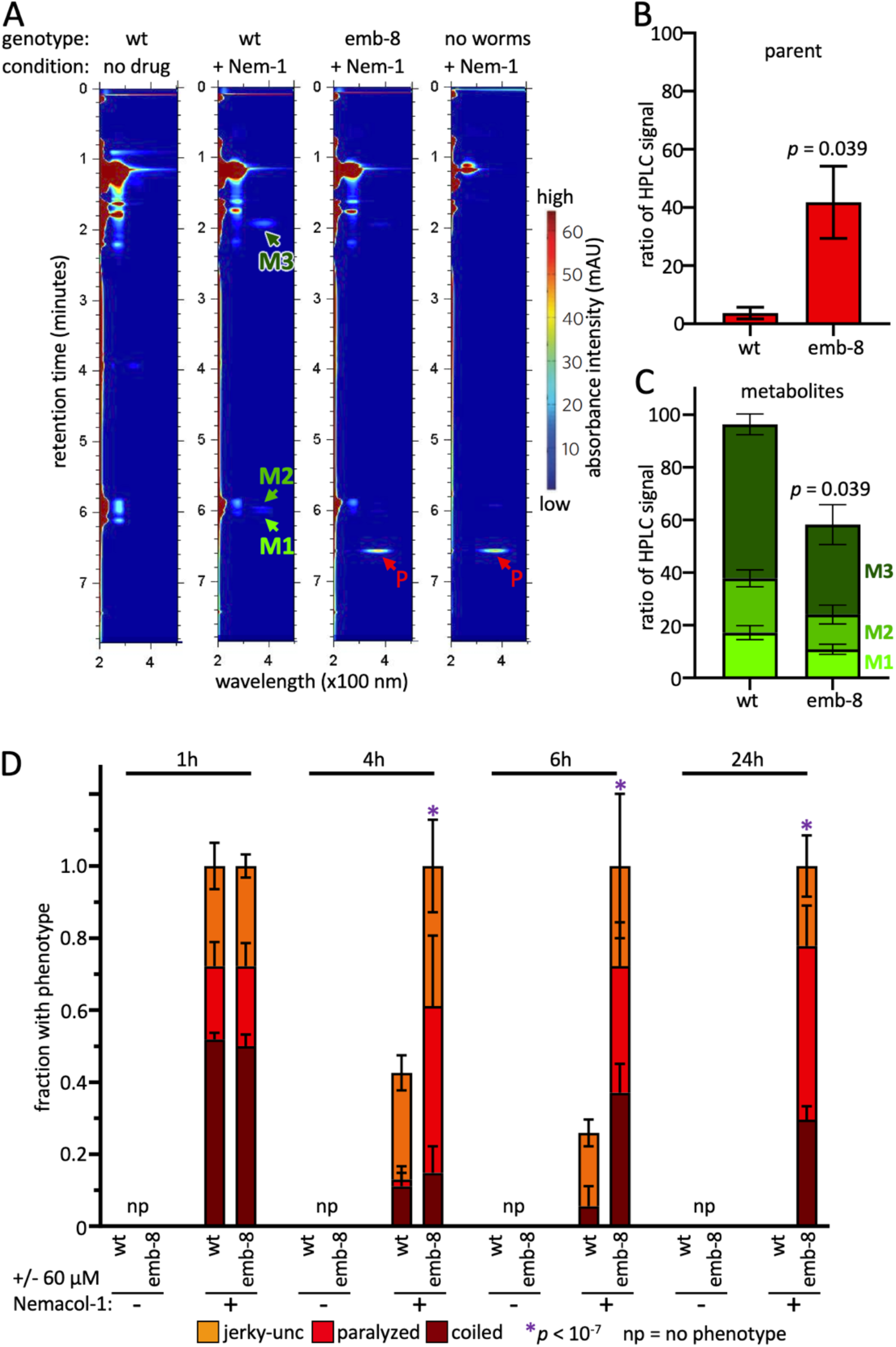
*C. elegans* animals with defective cytochrome p450 reductase fail to metabolize Nemacol-1 and fail to recover to Nemacol-1 over time. (**A**) Heat-mapped chromatograms of exemplar HPLC runs that are coupled to a diode array detector (HPLC-DAD) of wild-type and EMB-8-knockdown animals treated with solvent-only control or 60 μM Nemacol-1. Retention time is shown on the y axis, and absorbance wavelength is shown on the x axis. The scale of absorbance intensity, in milli–absorbance units (mAU), is shown on the right. The parent Nemacol-1 peak (red arrow), and Nemacol metabolite peaks (green arrows) are indicated. See methods for HPLC-DAD methodology. (**B**,**C**) Quantification of the relative ratio of parent and metabolite peaks to the total compound related signal from triplicate measurement of 2000 adult worms by HPLC-DAD (see methods). Reporting the mean with the standard error of the mean. *P* values were generated using unpaired two-sided students t-test. (**D**) EMB-8 disruption suppresses the dissipation of the Nemacol-induced phenptypes. L4 worms were picked onto solid agar containing 60 μM of molecule and scored at the indicated concentration and indicated time. The indicated phenotypes are scored based on subjective classification (see methods for details). Data are the mean of 3 biological replicates scoring ∼18 animals per trial. * p < 0.001 using Chi-square test comparing the fraction of animals exhibiting any motor phenotype between WT and the indicated test mutant.

**Figure S2.**
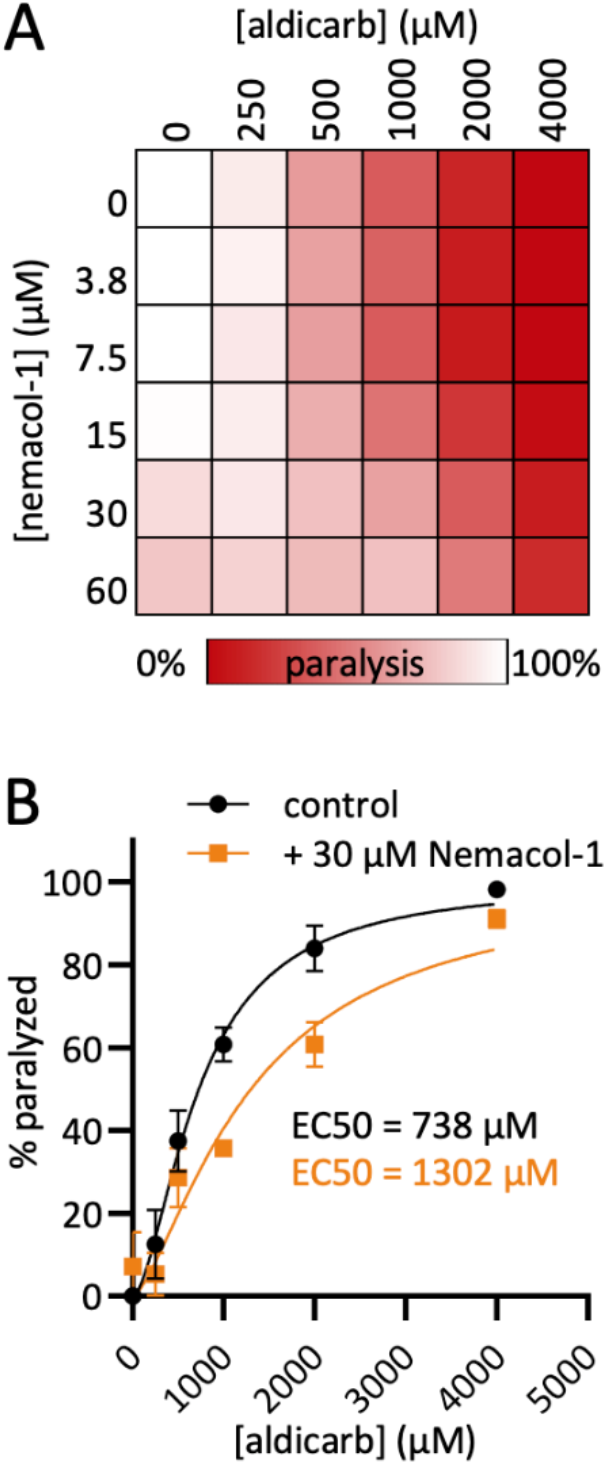
Nemacol-1 suppresses paralysis induced by the carbamate acetylcholinesterase inhibitor aldicarb. (**A**) Double dose-response matrices of Nemacol-1 + aldicarb showing the fraction of animals that were scored as paralyzed after 80 minutes. The benchmark for paralysis was a lack of sinusoidal body posture and an inability to back at least half a body length upon a touch on the head with a platinum wire. Values in cells represent the percent of animals scored as paralyzed. Data are the mean of 3 biological replicates scoring 28 animals per condition. (**B**) Dose-response curves from A highlighting the shift of aldicarb + 30 μM Nemacol-1. For control compared to 30 μM Nemacol-1 *p* = 9.8 × 10^−5^. P-value calculated using an extra sum-of-squares F test comparing the aldicarb dose response curves +/- 30 μM Nemacol-1 in GraphPad Prism version 9.3.1.

**Figure S3.**
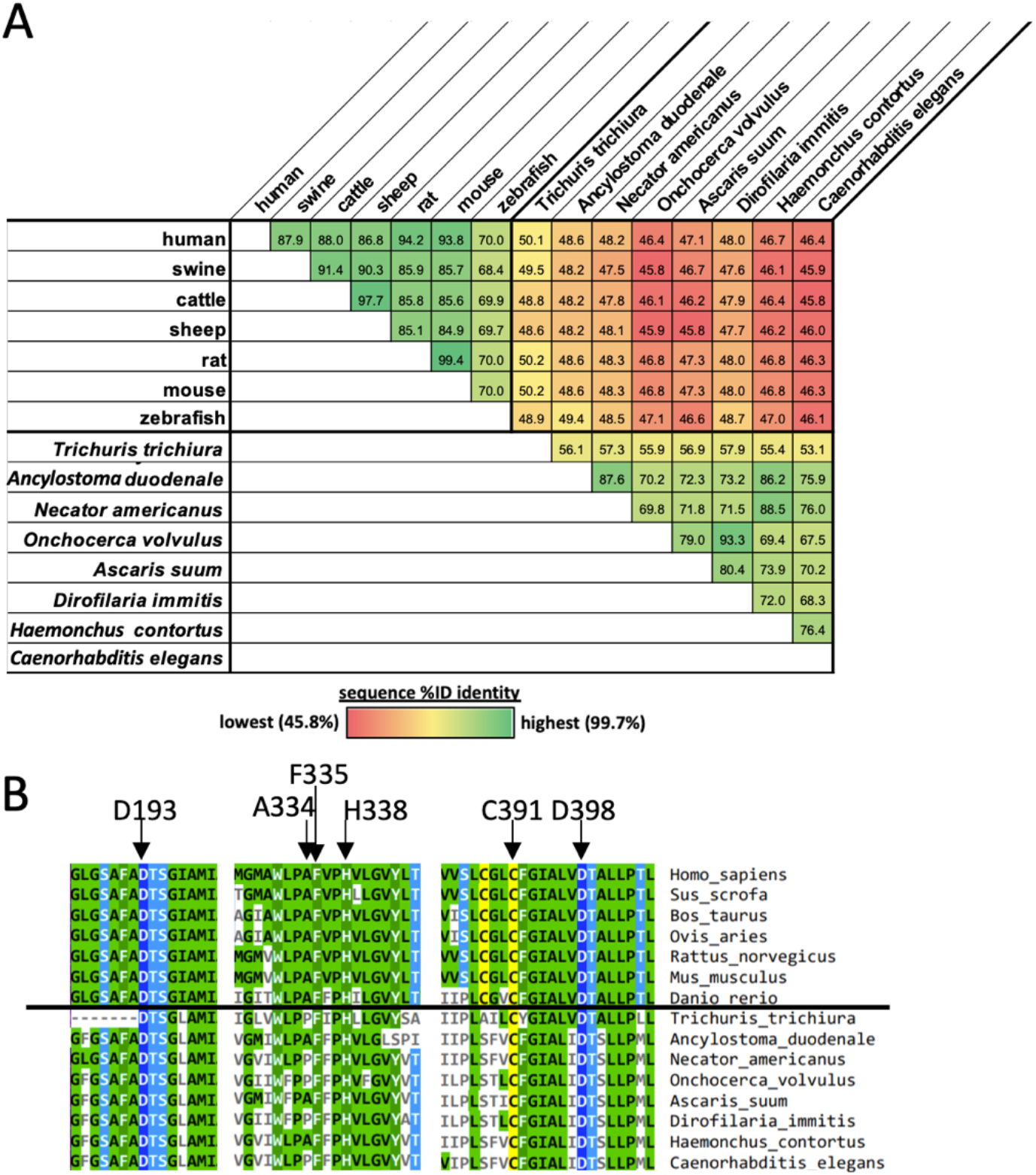
VAChT sequence differences exist between nematode and vertebrates. (**A**) VAChT sequence identity matrix comparing the percentage of residues that are shared between each species VAChT sequence. Sequences were identified as the closest homolog to human VAChT identified through BLAST searches^67^. Sequences used (NCBI reference sequence identifiers or otherwise stated): human NP_003046.2; swine (*Sus scrofa*): XP_013838900.2; cattle (*Bos taurus*): XP_002699016.1; sheep (*Ovis aries*): XP_027818269; mouse (*Mus Musculus):* NP_068358.2; rat (*Rattus norvegicus):* NP_113851.1; zebrafish (*Danio rerio*): NP_001071018.1; *Trichuris trichiura:* CDW52212.1; *Ancylostoma duodenale:* KIH66835.1; *Necator americanus:* XP_013297134.1; *Onchocerca volvulus:* A0A2K6VZC1 (UniProt ID) *Ascaris suum:* AgB02_g088_t01 (WormBase ParaSite transcript ID); *Dirofilaria immitis:* nDi.2.2.2.t09212 (WormBase ParaSite transcript ID); *Haemonchus contortus:* A0A7I4YIM0 (UniProt ID); *C. elegans:* NP_001379838.1. (**B**) Multiple sequence alignment of VAChT sequences shown in panel A highlighting residues that have been reported to decrease vesamicol affinity when mutated^31,36^. VAChT sequences are separated by vertebrate sequences (top) and nematode sequences (bottom). Showing 7 residues flanking vesamicol specificity determinants. Residue numbers are the position in the human sequence (D193 & D398^68^, A334^35^, F335^37^, H338^38^ and C391^36^).

